# In-field bioreactors demonstrate dynamic shifts in microbial communities in response to geochemical perturbations

**DOI:** 10.1101/2020.04.16.044594

**Authors:** Regina L. Wilpiszeski, Caitlin M. Gionfriddo, Ann M. Wymore, Ji-Won Moon, Kenneth A. Lowe, Mircea Podar, Sa’ad Rafie, Matthew W. Fields, Terry C. Hazen, Xiaoxuan Ge, Farris Poole, Michael W.W. Adams, Romy Chakraborty, Yupeng Fan, Joy D. Van Nostrand, Jizhong Zhou, Adam P. Arkin, Dwayne A. Elias

**Affiliations:** Oak Ridge National Laboratory, Oak Ridge, Tennessee, USA; University of Tennessee, Knoxville, Tennessee, USA; Montana State University, Bozeman, Montana, USA; University of Missouri, Columbia, Missouri, USA; Institute for Systems Biology, Seattle, WA, USA; University of Washington, Seattle, WA, USA; University of Georgia, Athens, GA, USA; Lawrence Berkeley National Lab, Berkeley, CA, USA; University of Oklahoma, Norman, OK, USA; University of California at Berkeley, Berkeley, CA, USA

**Author notes:** This manuscript has been authored by UT-Battelle, LLC under Contract No. DE-AC05-00OR22725 with the U.S. Department of Energy. The United States Government retains and the publisher, by accepting the article for publication, acknowledges that the United States Government retains a non-exclusive, paid-up, irrevocable, world-wide license to publish or reproduce the published form of this manuscript, or allow others to do so, for United States Government purposes. The Department of Energy will provide public access to these results of federally sponsored research in accordance with the DOE Public Access Plan (http://energy.gov/downloads/doe-public-access-plan). These authors contributed equally to this work (Regina L. Wilpiszeski, Caitlin M. Gionfriddo).

## Abstract

Subsurface microbial communities mediate the transformation and fate of redox sensitive materials including organic matter, metals and radionuclides. Few studies have explored how changing geochemical conditions influence the composition of groundwater microbial communities over time. We temporally monitored alterations in abiotic forces on microbial community structure using 1L in-field bioreactors receiving background and contaminated groundwater at the Oak Ridge Reservation, TN. Planktonic and biofilm microbial communities were initialized with background water for 4 days to establish communities in triplicate control reactors and triplicate test reactors and then fed filtered water for 14 days. On day 18, three reactors were switched to receive filtered groundwater from a contaminated well, enriched in total dissolved solids relative to the background site, particularly chloride, nitrate, uranium, and sulfate. Biological and geochemical data were collected throughout the experiment, including planktonic and biofilm DNA for 16S rRNA amplicon sequencing, cell counts, total protein, anions, cations, trace metals, organic acids, bicarbonate, pH, Eh, DO, and conductivity. We observed significant shifts in both planktonic and biofilm microbial communities receiving contaminated water. This included a loss of rare taxa, especially amongst members of the *Bacteroidetes*, *Acidobacteria*, *Chloroflexi*, and *Betaproteobacteria*, but enrichment in the Fe- and nitrate-reducing *Ferribacterium* and parasitic *Bdellovibrio*. These shifted communities were more similar to the contaminated well community, suggesting that geochemical forces substantially influence microbial community diversity and structure. These influences can only be captured through such comprehensive temporal studies, which also enable more robust and accurate predictive models to be developed.

## INTRODUCTION

Microorganisms are known to substantially contribute to the fate of redox sensitive contaminants in the subsurface ^1–5^. However, subsurface microbial community structure and function can be impacted by various physicochemical, geological, and biological conditions including temperature, redox potential, groundwater flow, influx of contaminants, chemical toxicity, available nutrients and carbon sources, and microbial competition ^6–11^. It is unclear whether the subsurface biotic potential to transform contaminants is driven primarily by the resident geochemistry or by indigenous microbial community structure. Thus, a fundamental question arises as to whether similar *in-situ* geochemical conditions will drive different microbial communities towards similar community structures. Field studies exploring 16S rRNA gene diversity across contaminant gradients show clustering of biomarkers with geochemical variables ^12^ with nitrate being one strong geochemical factor correlating with ecological association ^13^.

The Bear Creek aquifer is located within the Oak Ridge Reservation (ORR; Oak Ridge, TN) and consists of 243 acres of contaminated, and 402 acres of uncontaminated, areas ^14^ underlying interbedded shales, siltstones, and limestones of the Conasauga Group ^14^. The site contains a gradient of contaminants generated during the research and production of nuclear materials, including nitrate, sulfate, various metals (e.g., Cr, Ni, Cu), radionuclides (e.g., U and Tc), and organic contaminants (i.e., tetrachloroethylene and toluene) ^15^ leaching from unlined disposal ponds that were used until 1983 ^15^, with pH ranging from 3-8 ^14^. The complex nature of the contaminant plumes and their influence on microbial community diversity and structure, as well as the biotransformation potential of the *in-situ* community, is largely unknown. While there have been several studies conducted at the ORR ^11–13, 16–24^, most were conducted on groundwater wells with sporadic or periodic sampling to provide a series of snapshots. These studies included both geochemical and microbial community diversity data but could not capture the real-time or near real-time biochemical responses to contaminant stress.

Throughout the numerous snapshot studies at the ORR, several abundant organisms have been characterized including *Azoarcus* spp. from the acidic contaminated area as well as *Rhizobium* spp. and *Diaphorobacter* spp. from the nitrate-elevated circumneutral pH area which have been implicated in nitrogen-fixation or nitrate-reduction^12^. In the uncontaminated area, abundant organisms have included the sulfur-oxidizing *Thiobacillus thioparus*, ammonium-(*Nitrosomonas europaea*), nitrite-(*Nitrospira marina*), and iron-(*Acidovorax ebreus*) oxidizing organisms as well as a number of *Acidobacteria*, *Actinobacteria*, *Bacteroidetes*, *Chloroflexi*, *Deinococcus*-*Thermus*, *Firmicutes*, *Nitrospira* and α-, β-, γ-, δ-, and ε-*Proteobacteria*, comprising a diverse microbial community ^25^. Many of these organisms participate in N-cycling and play an important role in remediation ^26^ both in general and at the ORR^27^, and injection of carbon sources into the aquifer accentuated NO_3_ -utilization, accumulation of NO_2_ -, and production of N_2_O ^28^.

The question at hand is to what extent does the local geochemistry shape the microbial community structure, and therefore function? To address this, we investigated the structure and development of the local microbial community through the use of in-field bioreactors. These in-field bioreactors enable us to operate at the groundwater source indefinitely ^25^ allowing for constant monitoring of the microbial community and its responses to perturbation in real-time while allowing for assessment of both planktonic and biofilm communities (eg^29^). The goal of this study was to begin with the local background microbial community, allow it to self-stabilize in the bioreactors and then induce a multiscale perturbation by titrating in the contaminated groundwater, analogous to contaminant plume movement. By doing so, we could assess whether the introduction of the contaminated groundwater shaped the established microbial community into a less diverse community that was more similar to the *in-situ* community from the contaminated groundwater.

## MATERIALS AND METHODS

### Bioreactor experimental set-up

The ORR site itself is comprised of four main areas: background, Area 1, Area 2 and Area 3. The local geological and microbial features of the ORR background study site has been well described previously ^13–15^. We established small, in-field bioreactors located within a mobile laboratory in the background area to obtain and study *in-situ* groundwater and the associated microbial communities directly from an established well ^25, 29^. Groundwater was pumped from well FW305 into a sterile, 1L source reservoir container at a rate of 100 mL min^−1^, resulting in a residence time of 8 minutes to minimize any “bottle effect” (Figure S1A). The source reservoir and bioreactors were continuously flushed with 13% O_2_/87% N_2_ gas (Airgas, Radnor PA) to supply the bioreactors with *in-situ* dissolved oxygen levels of 2.2-6.3 ppm O_2_ as previously described ^25, 29^.

Each bioreactor had an external water jacket to maintain the bioreactor at ~14°C, the typical *in-situ* groundwater temperature. Groundwater was transferred through peristaltic pumps and purged more than 48 hours to stabilize a homogeneous groundwater supply before beginning flow to the in-field bioreactors at ~0.22 mL/min. Each bioreactor was filled with 800 mL groundwater and 100 mL of headspace, with eight removable biofilm coupons (Figure S1B). Core sediment (FW305-05-36) collected during the drilling of groundwater well FW305 and stored at −80oC was homogenized via freezer-mill (Freezer/Mill® 6870, SPEX® Sample Prep, Metuchen, NJ) at 25g/run with a sequence of 2 min freeze, 3 cycles of 2 min grind, and a 1 min pause for cooling. Samples were freeze dried, homogenized again, and sieved (500 µm) for uniformity. Each coupon was filled with 2±0.005 g of freeze-dried and sieved sediment, then assembled into in-field reactors and autoclaved. A total of 8 coupons and 1 stir bar were placed into each reactor. An additional sediment aliquot was ground and examined using an X-ray diffractometer (XRD, X’pert PRO, PANalytical, Natick, MA). The detailed specification and preparation of each custom-designed in-field bioreactor has been previously reported including assemblage, sterilization, gas purging, groundwater dripping through a tube breaker, and stirring ^25^. For sample harvesting the coupons were withdrawn from the reactor, placed in individual sterile bags and kept on ice until the coupons could be disassembled. Each coupon sediment was transferred to a sterile, DNase free tube and stored at −80oC prior to analysis ^30^.

Six in-field bioreactors were set up and operated simultaneously: three test bioreactors, and three control bioreactors that did not receive the contaminated groundwater. All reactors were slowly filled with unfiltered groundwater from well FW305 (day 0) until the sediments in the coupons were submerged, and the reactors were completely filled to 800 mL after 24 hours. After three days of acclimation to establish a microbial community, groundwater flow was started (day 4) and the reactors were sampled twice weekly (Monday/Thursday). For the first two weeks filtered FW305 groundwater flowed into each bioreactor through inline 0.2 µm filters (6 drops/min, 16.7 mL/hr, 0.3 mL/min; each drop was 0.05 mL) that were changed every two days to provide homogeneous dissolved geochemical condition and saved for gDNA isolation. After bioreactor sampling on day 18, four of the eight biofilm coupons from each reactor were collected to serve as reference of the FW305 microbial community and were replaced with new sterile coupons. The test bioreactors were then switched to receive filtered nitrate-elevated groundwater from well FW706 with the same flow rate. To minimize geochemical variation, groundwater from FW706 was all collected on a single day, filtered (0.22 μm), dispensed into 2-liter culture bottles without headspace and stored at 4oC. Bottles were replaced when the previous bottle contained <10% volume, and the same groundwater characterizations were performed on each new bottle.

Liquid samples were collected at the outport of the bioreactors with a 60 mL syringe. For each planktonic sampling, ~60 mL were collected including 1 mL for protein analysis, 1 mL preserved in 10% formaldehyde for cell counts, ~2 mL stored in gas-tight vials without headspace for bicarbonate titration, ~5 mL for metal analysis, and ~5 mL for physico/geochemical measurements including pH, conductivity, and dissolved oxygen (DO) measured immediately upon acquisition in the mobile laboratory. An additional 8 mL subsample was immediately dispensed to N_2_-flushed anoxic pressure tubes using a needle and syringe for later oxidation-reduction potential (ORP) measurement in an anaerobic chamber (Coy Laboratory Products, Inc., Ann Arbor, MI)^15^. The remaining 40 mL of planktonic sample was filtered through 0.22 µm nitrocellulose in a polypropylene filter holder (Swinnex, Germany). The first 15 mL of filtrate was dispensed into a 15 mL falcon tube, sub-aliquoted, and stored at 4oC until analysis for organic acids and anions via ion chromatography (Dionex). 2 mL of filtrate were frozen for metal analysis, and an additional 1 mL was acidified in-field with 100 µL 1M HCl and stored at 4oC prior to analysis for cations via ion chromatography. Community gDNA was obtained from the filters and all biofilm coupon sediments collected on day 18 and at the completion of the experiment and sequenced for 16S rRNA.

### Characterization of groundwater and reactor water

The pH, conductivity, and DO were measured twice weekly (Monday/Thursday) with an electrochemistry benchtop meter, Orion^TM^ Versa Star Pro^TM^ (Thermo Fisher Scientific, Waltham, MA), combined with Ross^TM^ Ultra pH/ATC Triode (Orion^TM^ 8157 BNUMD), conductivity cell (Orion^TM^ 013005MD), and DO probe (Orion^TM^ 083005MD), respectively. ORP was measured in an anaerobic glove bag using a combination of a benchtop meter, Star logR and a Triode ORP probe, Orion 9180BN (Thermo Scientific). The bicarbonate concentration was measured by titration to pH 4.5 with 10 mM HCl using a Dosimat 765 (Metrohm, Switzerland). Cell counts were performed by acridine orange direct counting. The protein analysis used a bicinchoninic acid protein assay kit (BCA^TM^, Pierce, Rockford, IL) with bovine serum albumin as the standard ^31^.

The organic acids (lactate, acetate, propionate, formate, pyruvate, fumarate, succinate and citrate) (0.5‒200 μM) and anions (fluoride, chloride, nitrite, bromide, nitrate, sulfate, and phosphate) (1.0‒2000 μM) were identified and quantified with a Dionex^TM^ ICS-5000^+^ DP dual pump, ICS-5000^+^ DC dual column equipped with an AS11-HC column at 35oC with a KOH gradient of 1‒60 mM with a ICS-5000^+^ EG effluent generator system (Thermo Fisher Scientific). Samples volumes of 10 μl were injected with a Dionex^TM^ AS-AD auto-sampler at room temperature. The cations (lithium, sodium, ammonium, potassium, magnesium, and calcium) were identified and quantified with a Dionex™ ICS 5000^+^ series with a CS12-A column at 35 oC with an isocratic 20 mM methanesulfonic acid effluent at 1 mL/min. Samples were acidified with a 10 vol.% 1M HCl solution before analysis. The detection range for cation measurements was 10‒5000 μM.

Trace metal concentrations were determined based on EPA Method 6020A. Both filtered and unfiltered water samples were vortexed and diluted 1:5.75 into 2% (vol/wt) trace-grade nitric acid (VWR, Radnor, PA) in acid-washed polypropylene tubes and stored overnight at ambient temperature. The diluted samples were pelleted and the clarified samples were transferred to another acid-washed polypropylene tube for measurement of 16 (Al, K, V, Cr, Mn, Fe, Co, Ni, Cu, Zn, As, Mo, Cd, W, Pb, and U) elements using an Agilent 7900 Inductively Coupled Plasma Mass Spectrometer (ICP-MS) fitted with MicroMist nebulizer, UHMI-spray chamber, Pt cones and an Octopole Reaction System (ORS) collision cell (Agilent Technologies, Santa Clara, CA). The data was collected using a 3-point peak pattern, 3 replicates, 100 sweeps/rep and various integration times (all >0.3 s) using either no gas or He gas in the ORS collision cell. The external calibration standards, IV-ICPMS-71A and IV-ICPMS-71B, were used to create an 11-point curve from 0 - 500 ppb for each element and the internal calibration standard, IV-ICPMS-71D, was added to each sample (Inorganic Ventures, Christiansburg, Virginia). External calibrations were performed every 132 samples or each day, whichever occurred first. Data was processed using the nearest internal standard by mass and fitted to a linear curve though the calibration blank. Three elements (Na, Mg, and Ca) were measured by IC analysis.

### Characterization of soil samples

The mineral composition of pulverized sediment in the biofilm coupons was characterized with an XRD equipped with Mo-Kα radiation at 55 kV/40 mA between 5‒35° 2θ with 1.5° 2θ/min ^32^ Soil pH was measured with core sediment reacted with a 10 mM CaCl_2_ solution (soil:solution ratio of 1:2 g/l) equilibrated for 60 min.

Samples for carbon and nitrogen analysis were ground to a fine powder, oven-dried at 70°C and stored in a desiccator. Approximately 0.2 g of prepared sample was analyzed on a LECO TruSpec elemental analyzer (LECO Corporation, St. Joseph, MI). Triplicate standard of known carbon and nitrogen concentration (Soil lot 1010, LECO Corporation, carbon = 2.77 % ± 0.06 % SD, nitrogen = 0.233 % ± 0.013 % SD) were used to ensure the accuracy and precision of the data.

### DNA extraction and microbial community sequencing

Microbial gDNA was extracted from biofilm sediment coupons using the DNeasy PowerMax Soil Kit (QIAGEN, Germantown, MD USA). Microbial genomic DNA was extracted from nitrocellulose and acrodisc filters using the DNeasy PowerSoil DNA isolation Kit (QIAGEN, Germantown, MD USA) following the manufacturer’s instructions. DNA quantity was determined using PicoGreen ^33^.

A two-step PCR method was used for library preparation as described previously^34^. Briefly, primers 515F and 806R were used in the first step PCR to amplify the V4 region of prokaryotic 16S rRNA genes (515F [5’-GTGCCAGCMGCCGCGGTAA-3’] and 806R [5’-GGACTACHVGGGTWT CTAAT-3’]). All amplifications were performed in triplicate and then pooled and purified using Agencourt AMPure XP magnetic beads. Phasing primers, designed with 0-7 base-length spacers to increase base diversity, containing barcodes for sample identification were used for the second PCR. Both PCR amplifications used the following program [1 min at 94 °C followed by 10 (first step) or 20 cycles (second step) of 20 s 94 °C, 25 s at 53 °C, and 45 s at 68 °C, then a final 10 min extension at 68 °C]. Equal amounts (100 ng) of the final PCR product from each sample were pooled (approximately 130 samples per sequencing run). Sample libraries were then sequenced on a Miseq platform (Illumina, San Diego, CA, USA) at the Institute for Environmental Genomics Consolidated Core Laboratory (University of Oklahoma, Norman, OK, USA) as described previously^35^. The raw paired-end 16S rRNA amplicon sequencing files are available from NCBI under the BioProject ID PRJNA613078.

Raw sequences were demultiplexed, quality filtered, and trimmed using Trimmomatic, then forward and reverse reads were merged, dereplicated, and singletons removed using VSEARCH ^36, 37^. Chimeras were removed from dereplicated reads prior to OTU clustering using de novo and reference chimera filtering using VSEARCH and the SILVA gold reference database^37, 38^. The non-chimeric dereplicated nucleotide sequences were then clustered at 97% sequence identity and original trimmed reads were mapped to the centroid database using VSEARCH to produce OTU counts. Taxonomic classifications were assigned to centroid 16S rRNA sequences using the RDP-classifier at a confidence cut-off of 80%^39^. Downstream analyses, including statistics, of 16S rRNA amplicon sequencing data was performed using STAMP and the ‘phyloseq’ and ‘vegan’ packages in R ^40, 41^.

Random Forest classification and regression analyses were conducted using the sample-classifier module implemented in QIIME. Analyses were run using the 16S OTUs described above to fit models using 5-fold nested cross-validation with parameter tuning on and p-n estimator set at 100x. Each sample from each reactor at each time point was treated independently. Taxonomic identities for features significantly contributing to the models were assigned as described above.

## RESULTS AND DISCUSSION

### Physical properties of groundwater and bioreactor water

The source groundwater from the background site (well FW305) and groundwater from the nitrate- and heavy-metal contaminated site (well GW706) varied in pH, ORP, conductivity, and DO (Table 1). The pH of FW305 groundwater varied between 6.42–6.97 and the FW305-source reservoir between 6.73–7.10 throughout the experiment (Figure 1). The GW706 groundwater pH stored in 2-L glass bottles was 7.8‒7.97 throughout the experiment, slightly higher than the pH 7.52 measured at the well during groundwater collection. The pH in FW305 receiving bioreactors was 8.02–8.78, while bioreactors with GW706 water had slightly lower pH at 7.99–8.39 (Figure 1). The bioreactor pH was higher compared to source groundwaters, likely due to the removal of biogenic CO_2_ by stirring and/or dissolution of carbonate or saprolite from freeze-dried core sediment in the biofilm coupons^29^. We observed an increase in pH in the bioreactors post-coupon sampling (t = 18), possibly indicating that fine sediment released from coupons could be contributing to pH buffering. As sediments settled, a decrease in pH was observed between t=0 and t=18 (Figure 1). Despite differences in pH values between bioreactor and source groundwater, the temporal trends in the pH of the control bioreactors reflected the pH of the FW305-source reservoir (Figure 1).

**Table 1.** Comparison of groundwaterwell geochemistry as measured in-well.

|  | **FW-305**<br>**(Background)** | **GW-706**<br>**(Contaminated)** |
| --- | --- | --- |
| pH | 6.85 – 6.97 | 7.80 - 7.97 |
| DO (mg/l) | 2.3 - 3.8 | 2.2 - 3.1 |
| Temp | 16.7 | 14.2 |
| Cond (uS/cm) | 150 – 262 | 525 - 667 |
| NO<sub>3</sub> µM | 4.7 - 5.6 | 539 - 544 |
| SO<sub>4</sub> µM | 35 - 38 | 153 - 168 |
| DOC mg/L | 0.28 | 1.2 |
| Redox (mV) | -249 to -155 | -235 to -188 |
| Copper (µM ) | 0.001 - 0.006 | 0.010 - 0.062 |
| Nickel (µM ) | 0.005 - 0.020 | 0.025 - 0.048 |
| Uranium (µM ) | 0.002 - 0.003 | 0.189 - 0.233 |

**Figure 1.**
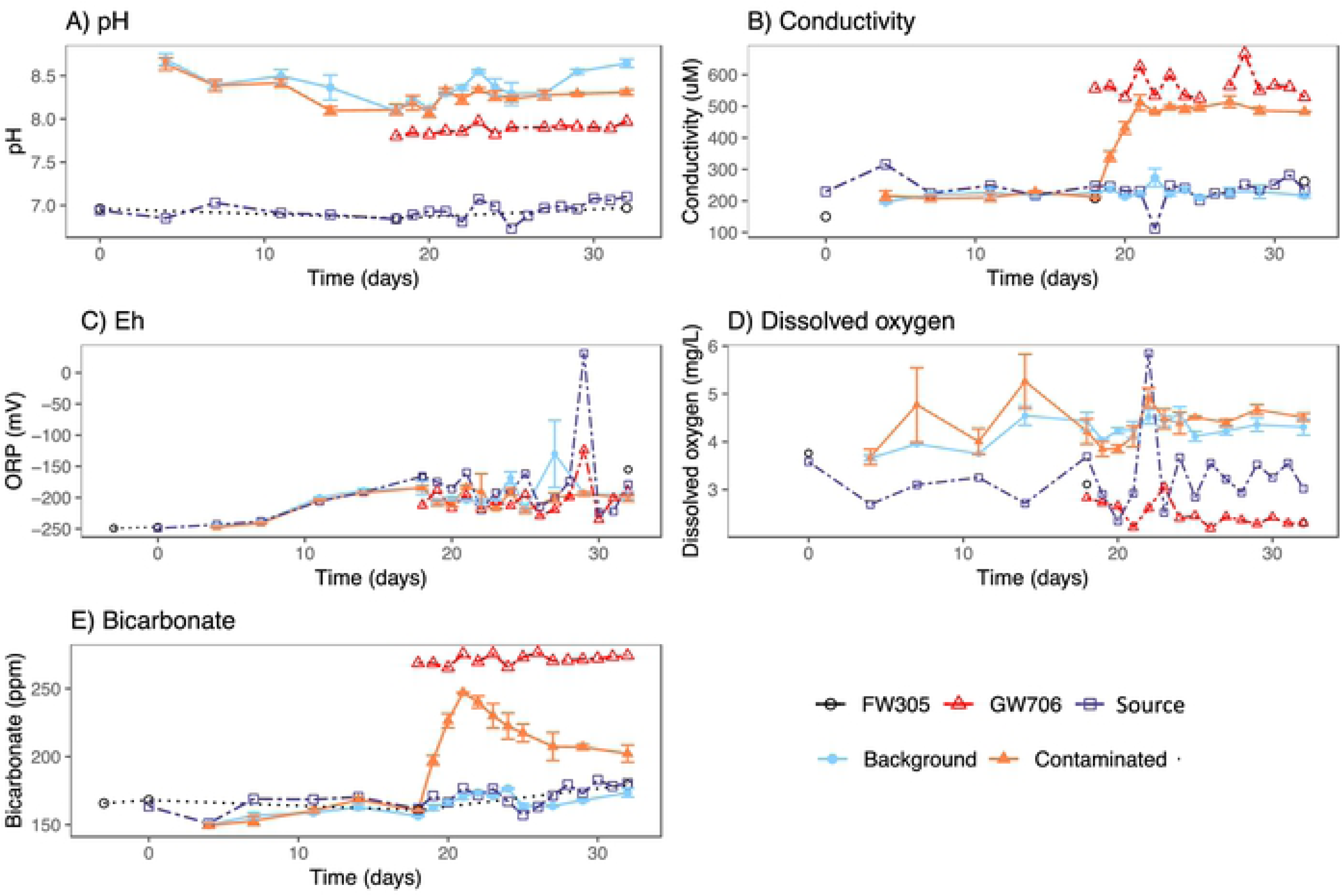
Physicochemical measurements from unfiltered water sampled from bioreactorsr receiving Background or Contaminated water, groundwater from wells FW305 and GW706, and the Source recervoir for the background reactors over the duration of experiment. Error bars represent one standard error.

Bicarbonate concentrations in FW305 water were 166–179 ppm while GW706 was higher at 266–276 ppm (Figure 1). The higher alkalinity of GW706 groundwater may reflect the geological setting of the two wells. Site GW706 is surrounded by interbedded limestone 42 and anthropogenic filling materials 15, compared to FW305 which contains severely weathered soil and saprolite 30. Bicarbonate concentrations in reactors reflected source groundwater concentrations, with higher bicarbonate in reactors receiving GW706 water (t=18) (Figure 1). The ORP increased in source and bioreactor waters from −250 mV to - 150 mV over the course of the experiment. Large fluctuations in FW305 can be attributed to rainfall events and the gradual decrease at the beginning of the experiment could be due to microbial community activity. Conductivity remained between 150‒262 µS/cm in FW305 groundwater and was similar in the FW305-source reservoir (112‒249 µS/cm) and FW305 bioreactors (187‒327 µS/cm) throughout the experiment (Figure 1). The GW706-reservoir conductivity was higher (525‒667 µS/cm), consistent with field measurements at the GW706 well, of 557 µS/cm. Reactors receiving GW706 water increased from 210‒213 µS/cm to 472‒559 µS/cm after switching on day 18 (Figure 1). The higher conductivity values reflect the GW706 groundwater geochemistry, which historically has exhibited elevated concentrations of total salts, nitrate (75.7 mg/L) and sulfate (28.2 mg/L) compared to FW305 nitrate (0.14 mg/L) and sulfate (6.80 mg/L) (monitored on December 2012, unpublished data).

There was a marked difference in dissolved oxygen (DO) in well FW305 (0.19 mg/L) and GW706 (0.09 mg/L) groundwaters measured down-well compared to the source reservoirs for FW305 (2.34 - 5.85 mg/L) and GW706 (2.19 - 3.05) throughout the experiment (Figure 1). DO measured in reactors receiving both FW305 water (2.30 - 6.32 mg/L) and GW706 water (3.66 - 5.21 mg/L) were quite similar to one another and to the reservoir values (Fig. 1). The spike in DO observed on day 22 (5.85 mg/L) and drop in conductivity (112 µS/cm) are indicative of a significant rainfall event that occurred on day 18 - 19. The greater DO levels within the bioreactors are likely caused by a combination of headspace purging with 13% O_2_-balanced N_2_, incomplete gas-tight sealing around the coupons, air permeation into the black Viton lines and incomplete gas-tight sealing of in-field bioreactors.

### Chemical properties of groundwater and bioreactor water

The cation profiles over time reflected the influent source water and presumed-dissolution from the sediment coupons (Figure 2). Magnesium (32-42 μM) was stable over time in FW305 groundwater, the FW305 source reservoir, and reactors receiving FW305 water. Potassium (61-83 μM), sodium (168-275 M), and calcium (240-360 μM) were more variable in the samples receiving water from FW305.

**Figure 2.**
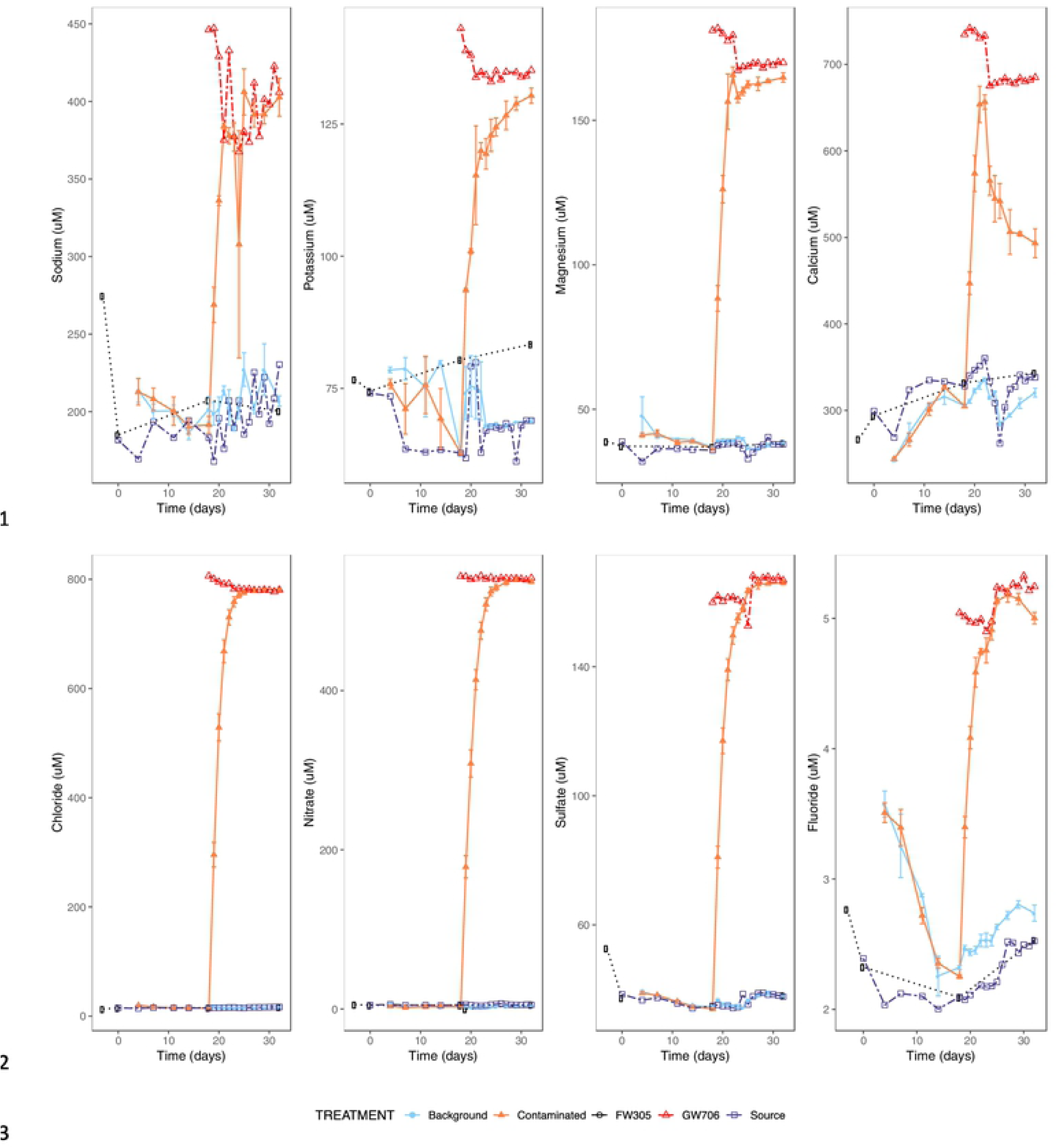
Major cation (top panel) and anion (bottom panel) concentration in filtered water collected from biorea1ctors receiving Background or Contaminated water, groundwater from wells FW305 and GW706, and the Source reservoir for the background reactors during duration of experiment. Error bars represent one standard error.

GW706 groundwater had consistently higher concentrations of these cations. Among the anions examined, fluoride (2-3.5 μM), chloride (13-22 μM), nitrate (2-7 μM), and sulfate (34-40 μM) were stable in FW305 groundwater, FW305-source reservoir and bioreactors receiving FW305 water. GW706 groundwater had consistently higher concentrations of anions, including F-(~5 μM), Cl-(~780 μM), NO -_3_ (~540 μM), and SO_4_^2^-(~165 μM). Anion concentrations in bioreactors receiving GW706 water increased to levels observed in GW706 water by day 24, six days following the switch from FW305 to GW706 water (Fig. 2).

Metal concentrations fell into 3 categories – those that rose in the bioreactors upon addition of GW706 contaminated water (eg uranium, potassium, copper, nickel), those that were relatively constant in reactors under both control and contaminated water (eg iron), and those that displayed a distinct peak- and-drop-off trend that coincided with introducing new sediment coupons (eg manganese) (Figure S2). Dissolved metals (those remaining after 0.2 μm filtration) accounted for most of the metal concentration except in the case of iron, for which particles > 0.2 μm accounted for up to 2-orders of magnitude greater concentration than the dissolved metal, consistent with the release of abiotic aggregates from the saprolite-rich sediment coupons. For manganese, the peak-and-drop-off pattern was exaggerated in the filtered (<0.2 μm) fraction of water from the bioreactors. This trend was consistent with leaching from the sediment coupons that were introduced at the start of the experiment and partially replaced when the water source was switched on day 18, with the concentration decreasing back to groundwater levels over 7 - 10 days.

In terms of organic acids, the FW305 groundwater has previously been shown to be oligotrophic ^25, 29^ with transient lactate and acetate peaks which decreased back to initial concentrations over time ^29^. In the current work, FW305 groundwater and control bioreactors exhibited <1 µM fluctuations in formate and acetate. Pyruvate, lactate, fumarate, propionate, succinate, citrate, and oxalate were all below the detection limit of 0.5 µM. However, the GW706 treated bioreactors showed wide fluctuations with increasing acetate and formate (Figure S3), suggesting greater dissimilatory microbial activity.

Total protein quantification as well as direct cell counts were used to measure temporal changes in planktonic microbial biomass. Overall, the trends in protein and cellular quantity were similar and unremarkable. Biomass steadily declined over the first two weeks of the bioreactor run then increased following the coupon change on t=18 (Figure S4). This increase was higher in the bioreactors receiving contaminated water compared to control bioreactors. Following the switch, biomass steadily declined in all reactors and remained at ~5 μg/mL protein and 500,000 cells/mL for the duration of experiment. Transient increases and decreases under both reactor conditions may be attributed to population-specific blooms within the community (see discussion below).

### Geological characteristics of sediment coupons

The homogenized sediment used in the biofilm coupons originated from the soil core recovered while drilling well FW305, the background groundwater source used in this study. The physical characteristics, including soil pH, mineral composition and carbon/nitrogen content of the core sediment used for establishing biofilm communities was characterized via XRD and elemental analyzer (Figure S5). The core sediment (FW305-06-06) had a pH of 5.39, which was slightly higher than previously reported (pH 4.78, FW305-05-24 ^30^ and pH 4.51, FW305-05-06 ^29^). The sediment was predominantly composed of weathered saprolite with little carbon and nitrogen (0.205 wt.% and 0.0152 wt. %, respectively). These values were similar to previous studies showing low total carbon content of 0.21‒0.225 wt.% which increased less than ~8 Δwt.% after carbon sequestration ^30^. Despite being the same mineral assemblage as core material used in previous studies ^29^, the present sediment aliquot had a higher XRD signal intensity suggesting a more mineralized layer with reduced amorphous or organic phases (Figure S5). A similar shift was observed previously as an increase in the relative intensity of clay mineral peaks after removing organics for gDNA extraction ^29^.

### Microbial community results

In the field, the natural microbial communities of the two groundwater sources used in this study reflected the different geochemical environments of the two water sources. The most abundant taxa were present in both wells (Figure 3) but field-collected planktonic samples from contaminated well GW706 were less diverse than the planktonic community observed in control well FW305 (Figure S6). Contaminated well GW706 was also significantly enriched in the xenobiotic genus *Phenylobacterium* which is known for using synthetic organic molecules like the herbicide chloridazon as a carbon source (Figure S7) ^43^.

**Figure 3.**
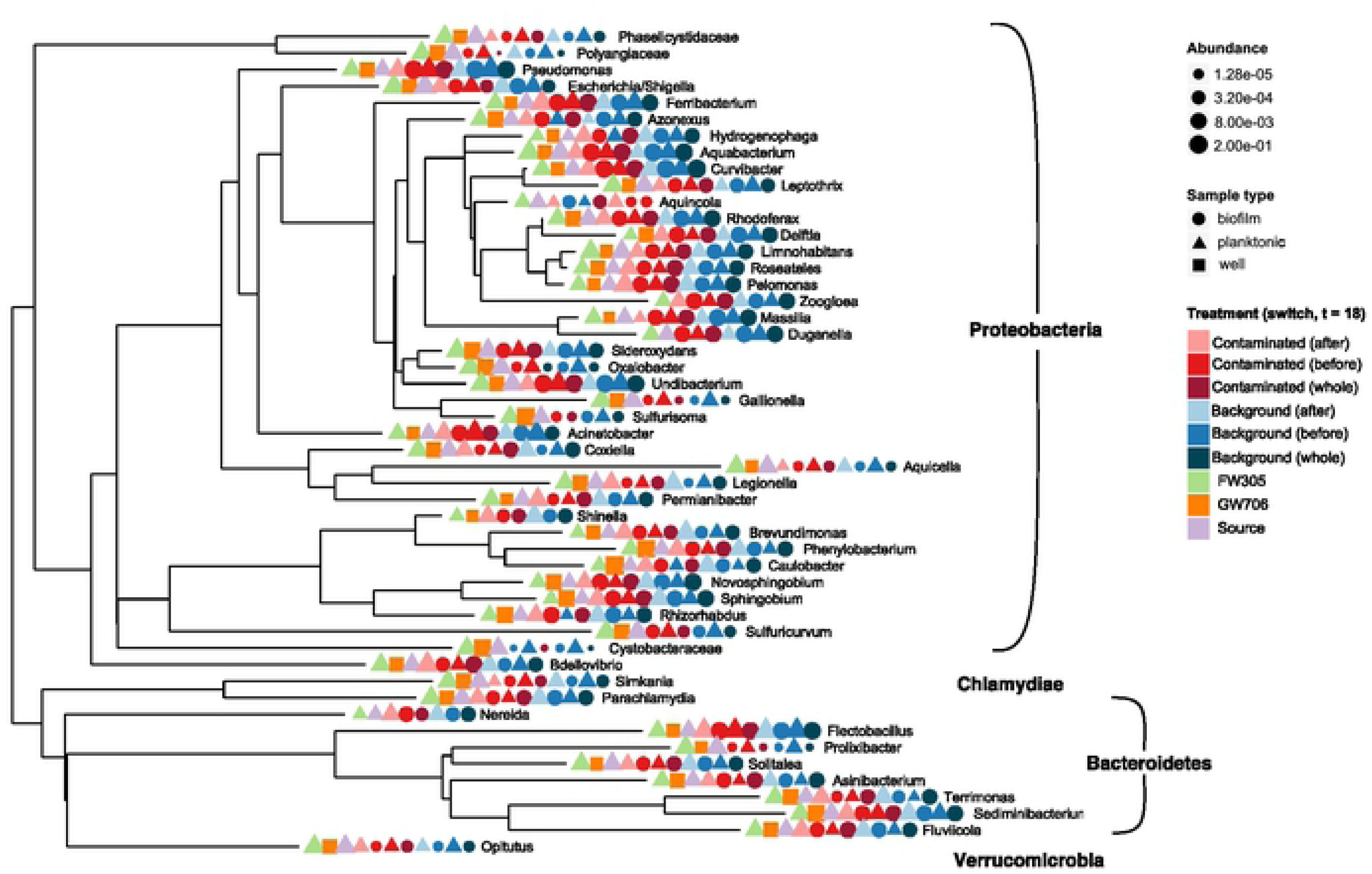
Phylogeny and relative abundance of top 50 (by abundance) 16S rRNA OTUs. OTUs have been merged at the genus level. Included in analyses are planktonic and biofilm communities from control (background, FW305 receiving) and contaminated (receiving GW706 water) reactor sampled before and after the switch from FW305 to GW706 water (day 18), and communitie from FW305 and GW706 wells and the source reservoir (FW305-receiving)

Within the bioreactors, the most abundant genera in both the planktonic and sediment-attached microbial communities generally matched the environmental water samples from the source wells (Figure 3) with notable differences in relative abundance for the attached community compared to the planktonic community. The initial stochasticity of the planktonic community during the first 18 days of the experiment shifted towards the established GW706 and FW305 well communities over time (Figure 4A).

**Figure 4.**
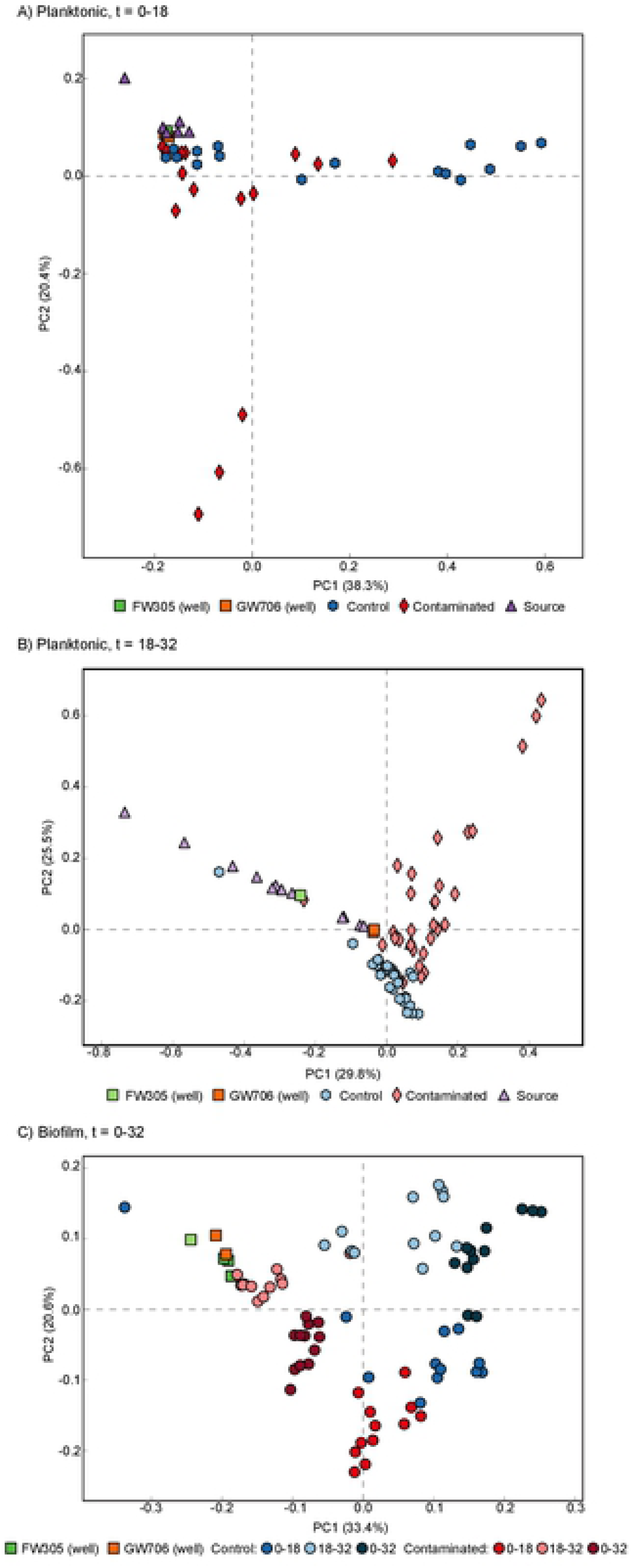
PCA plots of ANOVA distances showing variation in 16S rRNA genera abundance in A) planktonic communities from control (background, FW305 rec ivin) and contaminated (receiving GW706 water) reactors sampled before the switch on day 18; B) planktonic communities sampled after the switch to contaminated water (days 18-32); and C) biofilm communities collected at the start (days 0-18), end (days 18-32), or throughout the experiment (days 0-2).

The bioreactors also remained at a higher pH than the source water throughout the experiment (Figure 1). This trend was observed in previous experiments using these reactors ^25, 29^, and may reflect dissolution of calcium carbonate from clay minerals in the sediment-derived biofilm coupons (Figure S5). Despite this difference, the convergence of the bioreactor communities over the first 18 days suggests that pH was not the dominant parameter shaping these communities, and that the geochemical environment shaped and stabilized the establishing microbial community in the reactors.

After the water source was switched (t=18), the abundance of rare taxa decreased in the reactors receiving contaminated water (Figure S8). Those planktonic communities underwent a substantial shift in community composition while the reactors that continued to receive the control water changed very little (Figure 4B). The indigenous planktonic community in the source water feeding into the control reactors was highly variable over the latter half of the experiment, perhaps reflecting rain events, but the reactors themselves received filtered water and did not demonstrate such variability.

The biofilm communities showed a similar shift in composition after the switch to contaminated water. The community composition of coupons harvested from all reactors on day 18 were most similar to one another, while variability was most pronounced in biofilms established on day18 and harvested on day 32 (Figure 4C). The biofilm coupons that remained in reactors receiving contaminated water throughout the entire experiment shifted less than those coupons that were seeded when water was switched to GW706 on day 18. Similar trends were observed in the control reactor biofilms, although variability between sample replicates was more pronounced in control reactors than in those exposed to contaminated GW706 water. Overall, the community composition of the biofilms was less variable than the planktonic phase. This resilience of established biofilm communities to geochemical perturbations is consistent to previous work with in-field reactors^29^.

### Geochemical controls on microbial community composition

After the introduction of contaminated GW706 water on day 18, water in the in-field reactors reached comparable elevated concentrations of the major contaminants (nitrate, sulfate, chloride) ^12^ by day 23 with ~2.5 volumes of turnover. Multivariate analyses were used to assess how these geochemical constraints correlated with changes in the microbial community. The contribution of each independent factor could not be clearly determined since all contaminants were introduced from a single source, but environmental factors projected onto an ordination of the planktonic community showed significant (p ≤ 0.001) associations between the community composition and a number of covarying factors including nitrate, sulfate, conductivity, and various metals (Figure S10, Table S1).

Informed by the results of this multivariate analysis, a Random Forest algorithm was applied to create models fitting OTU data to categorical classifications (planktonic vs biofilm, control vs contaminated, before vs after switch) or to regressions against geochemical data (eg nitrate, sulfate, uranium). This type of machine-learning analysis has been applied to microbial community data sets previously^13, 44–46^. One advantage of machine-learning classification is the ability to distinguish features (such as OTUs) that best differentiate between categories or along geochemical gradients.

Classifying the community by experimental condition (contaminated groundwater, uncontaminated groundwater, source reactor, contaminated reactors, uncontaminated reactors) resulted in an overall model accuracy of 94.4% for biofilm samples and 85.5% for planktonic samples (Figure 5). Random Forest regressions of the community against select co-varying geochemical concentrations present at high concentrations in the contaminated water (nitrate, sulfate) provided a good fit for the planktonic community (R^2^ = 0.76, 0.81 respectively) but did less well for describing the biofilm community (R^2^ = 0.58, 0.66), which likely reflects the different time-scales at which planktonic vs biofilm communities respond to geochemical perturbations. Nitrate-reducing bacteria were of particular interest and were expected to dominate the microbial community in reactors fed with water from contaminated well GW706 based on previous measurements of decreasing nitrate concentration over time and the predominance of nitrate as a major contaminant in the ORR (Area 1 >199 mM, Area 2 >124 mM, and Area 3 >803 mM) ^15^.

**Figure 5.**
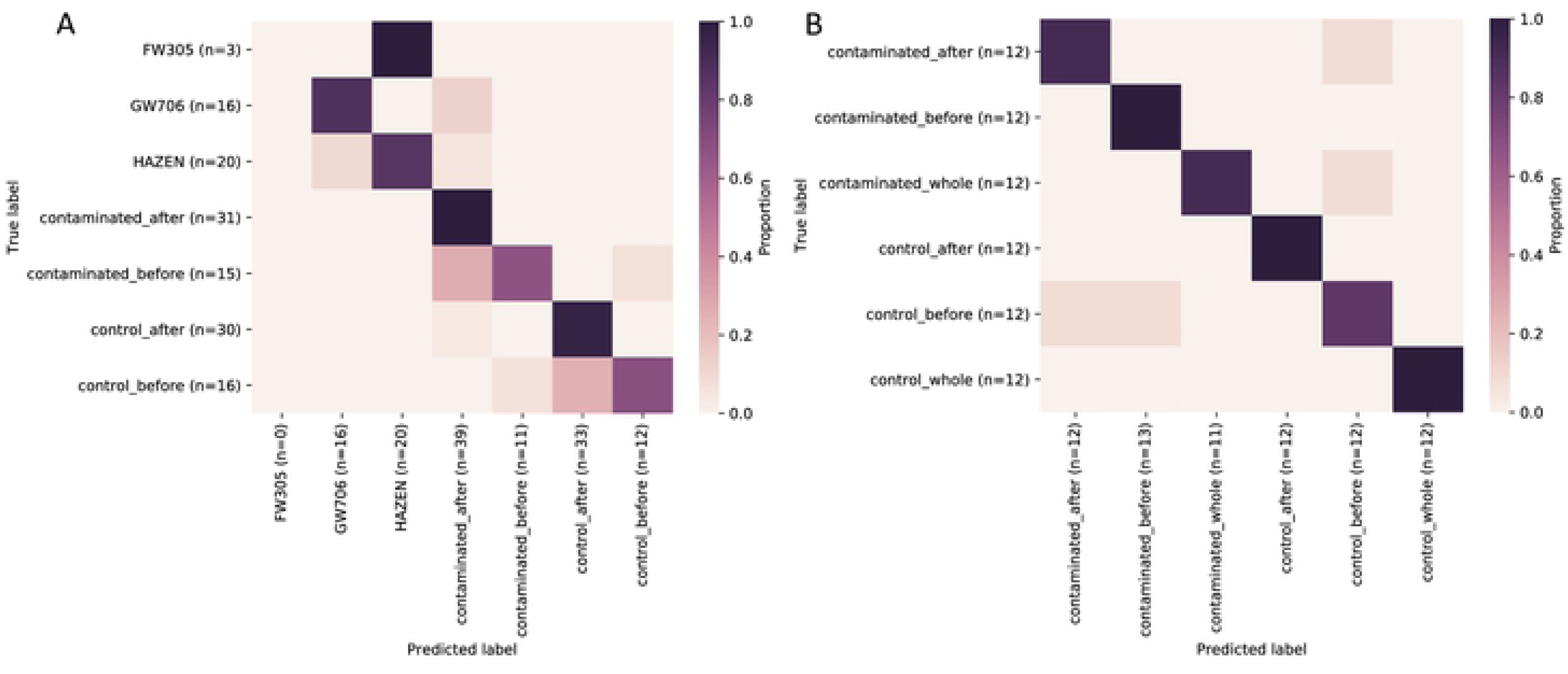
Random Forest models for planktonic (A) and (B) communities categorized by experimental condition. Overall model accuracy was 85.5% for planktonic samples and 94.4% for biofilm samples.

Within these models, the top-ranked taxa for correlating planktonic contaminant concentration to community composition included the genera *Massilia, Sideroxydans, Duganella, Brevinema, Sphingobium, Aquabacterium, Leptonema, Mesorhizobium*, and *Simkania*. For biofilm classification by category, the top-ranked taxa included *Curvibacter, Parcubacteria, Hydrogenophaga, Caulobacter*, and *Acinetobacter*. Several of these genera include nitrate-reducing or denitrifying members, many of which have been identified previously in 16S rRNA gene studies of nitrate- and uranium-contaminated groundwater from the Oak Ridge Reservation, including *Hydrogenophaga*, *Acinetobacter*, *Mesorhizobium*, and *Sphingobium*^47^. Nitrate and concomitant contaminants have been significantly linked to the presence of members of *Oxalobacteraceae* and *Acinetobacter* in nitrate and uranium contaminated groundwater in Russia^48^ and *Massilia* with nitrate concentration in previous in-field bioreactor experiments^25^.

### Key taxa in changing communities

Overall, planktonic diversity shifted after the introduction of contaminated water with a notable loss of rare taxa. After the switch to GW706 water we observed significant (Δabundance >1.0%, p<0.05) loss of *Flectobacillus*, *Sphingobium*, *Pseudomonas* and *Undibacterium* with significant enrichment in *Bdellovibrio*, *Brevundimonas*, *Ferribacterium*, *Fluviicola*, *Pelomonas*, and *Variovorax* (Figure 6; Figure S8; additional discussion in supplemental text). *Aminobacter* relative abundance also increased in contaminated reactors post-switch, but the increase was not significant due to variability between reactors (Figure S9). Notably, *Phenylobacterium* was not enriched in the planktonic phase of bioreactors that received contaminated water from GW706 (Figure S9) despite being the only genus that was significantly enriched in the natural GW706 community relative to groundwater from background well FW305 (Figure S7).

**Figure 6.**
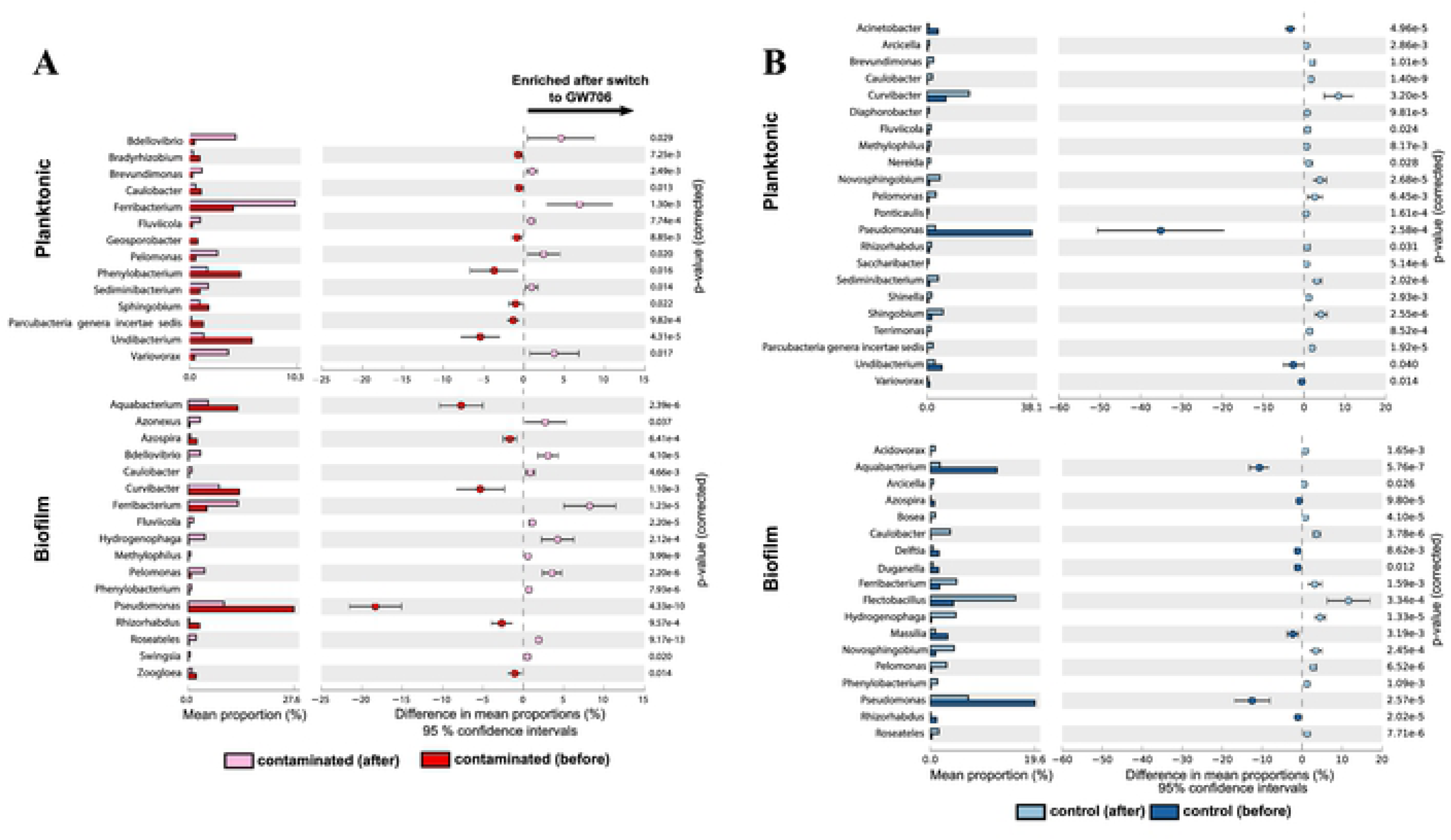
Significant differences (Fisher t-test, >0.5, p<0.05) in biofilm and planktonic microbial communities smapled before and after switch from FW305 to GW706 water (t =18) for A) reactors receiving contaminated water from GW706, and B) control reactors receiving only background water.

Within the biofilm communities, *Acinetobacter*, *Bdellovibrio*, *Caulobacter*, *Ferribacterium*, *Hydrogenophaga*, *Pelomonas*, *Phenylobacterium* and *Roseatales* were significantly enriched after the switch to GW706 water, while the relative abundance of *Aquabacterium*, *Azospira*, *Curvibacter*, *Pseudomonas*, *Rhizorhabdus* and *Zoogloea* significantly decreased (Figure 6A). In the control biofilm communities, *Caulobacter*, *Ferribacterium*, *Flectobacillus, Hydrogenophaga*, *Novosphingobium*, and *Pelomonas* were enriched after t=18, with a significant decrease in *Aquabacterium*, *Massilia*, and *Phenylobacterium* (Figure 6B). Compared to control reactors, the contaminated biofilm communities were enriched in *Acinetobacter*, *Azonexus*, *Bdellovibrio*, F*erribacterium*, *Pelomonas* and *Undibacterium* after t=18, while control reactor planktonic communities were dominated by *Caulobacter*, *Curvibacter*, *Flectobacillus*, and *Rhodoferax*.

Significant enrichment of the Fe(III)-reducing genus *Ferribacterium* and the parasitic genus *Bdellovibrio* occurred in both the planktonic and biofilm phases in contaminated water after the switch (Figure 6A). *Ferribacterium* reached up to 25% relative abundance of planktonic communities within the reactors that received contaminated water (Figure S9). This result was surprising given the consistency of iron between the control and contaminated water sources (Figure S2). However, members of the *Ferribacterium* have been seen in previous studies of contaminated water and sediment at the ORR^47, 49–51^ where they have been shown to reduce nitrate^52^ and possibly uranium (VI) ^50, 51^, both of which were significantly elevated in the contaminant water from GW706 (Figures 3, 4). *Bdellovibrio* species are motile, predatory *Deltaproteobacteria* that infect the periplasm of other bacteria^53^. This genus only became abundant (up to 30% relative abundance) in the reactors receiving contaminated water late in the experiment, and its peak coincided with a decrease in the more-abundant *Ferribacterium* and *Variovorax* (Figure S9). This may represent a predator-prey dynamic that could be investigated further for insight into predatory controls on nitrate-reducers in this environment.

This study provides a detailed view of abiotic forces on microbial community composition following the introduction of a contaminated groundwater source, analogous to contaminant plume movement. We observed significant shifts in both planktonic and biofilm microbial communities receiving contaminated water including a loss of rare taxa, especially amongst members of the *Bacteroidetes*, *Acidobacteria*, *Chloroflexi*, and *Betaproteobacteria*, with a simultaneous enrichment in the iron- and nitrate-reducing genus *Ferribacterium* and parasitic genus *Bdellovibrio*. The shifted communities showed periods of variability but ultimately converged, supporting a strong control on community structure by the geochemical environment.

## Acknowledgement

Authors appreciate the field support from Mehlhorn, T.L and Lowe, K.A. at Y-12 and background site. This material by ENIGMA-Ecosystems and Networks Integrated with Genes and Molecular Assemblies (http://enigma.lbl.gov), a Scientific Focus Area Program at Lawrence Berkeley National Laboratory is based upon work supported by the U.S. Department of Energy, Office of Science, Office of Biological & Environmental Research under contract number DE-AC02-05CH11231. Oak Ridge National Laboratory is managed by UT-Battelle, LLC, for the U.S. Department of Energy under contract DE-AC05-00OR22725. The funders had no role in study design, data collection and analysis, decision to publish, or preparation of the manuscript.

